# A structure-derived contact-network responsiveness atlas of human proteins

**DOI:** 10.64898/2026.06.22.733908

**Authors:** Ran Huang, Xin Ma, Dean Ta

## Abstract

Protein structures encode non-local contact organization, but static coordinates do not directly quantify how a contact network responds when effective stabilizing interactions are strengthened or weakened. Here we introduce **Contact-Network Responsiveness (CNR)**, a structure-derived framework that converts residue-level protein coordinates into density-controlled and topology-corrected response descriptors. The method is structure-source agnostic and can be applied to experimentally determined PDB structures, AlphaFold models, or other predicted structures; here, human AlphaFold models serve as the high-coverage structural substrate. Across 22,167 valid human protein structures, hydrophobic non-local contact density defined a nearly exact Bethe mean-field baseline for the conformational susceptibility threshold. A graph-aware residue-level extension then revealed systematic topology-dependent deviations from this density-only prediction. We define a topology correction ratio, 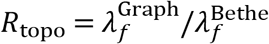, which separates topology-facilitated, density-dominated and topology-suppressed contact-network response regimes. CNR descriptors were associated with curated DisProt disorder annotations and broad-coverage UniProt/MobiDB-lite disorder fractions, supporting the interpretation that CNR captures a structural organization axis related to non-local contact availability and responsiveness.

## Introduction

Protein structures are usually interpreted as static coordinate models. Yet many biologically relevant perturbations do not act by changing the covalent chain itself, but by changing the effective strength, availability or organization of non-local contacts [1–3]. Ligand binding, mutation, changes in solvent quality, molecular crowding, ionic environment and post-translational modification can all alter how a protein contact network supports folded-like or contact-competent states. A structural model therefore contains more information than geometry alone, but this information must be converted into response descriptors before it can be compared across proteins.

This issue has become more important as high-coverage predicted structures enable proteome-scale structural annotation [4,5]. However, a static structure does not directly provide a response annotation. Confidence scores help identify reliable regions in predicted models, but they do not determine whether a protein’s non-local hydrophobic contact network is dense, easily activated, topology-suppressed, or disorder-associated. Because both experimentally determined PDB structures and predicted models contain residue-level coordinates, a general contact-network response framework should not be tied to a single structural source [4–6].

Here we introduce **Contact-Network Responsiveness (CNR)**, a structure-derived framework for converting protein coordinates into interpretable response descriptors. CNR first extracts non-local contact statistics from a protein structure, including the effective non-local coordination number *z*_nonlocal_, the hydrophobic contact fraction *r*_hydro_, and the hydrophobic non-local contact density

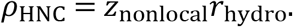

It then uses two response layers. A homogeneous Bethe mean-field baseline establishes the density-controlled response scale [13]. A graph-aware residue-level extension uses the actual hydrophobic contact graph to compute topology-dependent deviations from that baseline [1,14]. Together, these layers generate a CNR profile for each protein, including contact density, response thresholds, activation amplitude, susceptibility and topology correction.

CNR is not restricted to AlphaFold. It can be applied to experimentally determined PDB structures, AlphaFold models, or any predicted protein structure with residue-level coordinates. In this study, human AlphaFold structural models are used as a high-coverage substrate to generate a proteome-scale CNR atlas. We then test whether CNR descriptors are related to independent intrinsic-disorder annotations from DisProt and UniProt/MobiDB-lite [7–11]. This design separates three questions: how much response is explained by average hydrophobic contact density, how much is modified by contact topology, and whether low-contact or low-activation CNR regimes correspond to weak structural organization in external disorder annotations.

## Results

### CNR converts protein structures into contact-network responsiveness profiles

The CNR framework takes a protein structure as input and returns a multi-component contact-network response profile. Residues were represented by Cβ atoms, with glycine represented by Cα. Non-local contacts were extracted after excluding local sequence-neighbor contacts, and hydrophobic contacts were identified using residue hydrophobicity [1,2,12]. From these contacts, the first structural descriptor was the hydrophobic non-local contact density,

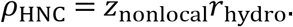

To make the CNR output explicit, each protein is represented by a descriptor profile rather than by a single scalar score (Fig. 2). The first component, *ρ*_HNC_, summarizes the average availability of hydrophobic non-local contacts in the structure (Fig. 2a). The Bethe layer then converts this density descriptor into a homogeneous response curve and defines the density-controlled threshold 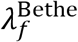 as the point of maximal susceptibility (Fig. 2b). The graph-aware layer uses the actual hydrophobic contact graph to compute residue-level activation, yielding the graph threshold 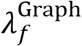, maximum activation 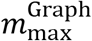, and maximum susceptibility 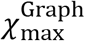 (Fig. 2c). Finally, the topology correction ratio

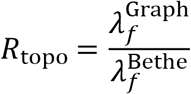

compares the graph-aware response with the density-controlled baseline, distinguishing topology-facilitated, density-dominated and topology-suppressed response regimes (Fig. 2d). The final CNR profile therefore combines density, threshold, topology correction, activation amplitude and cooperativity into a single structure-derived response annotation:

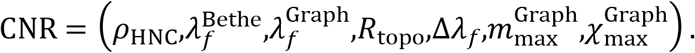

**Figure 1.**
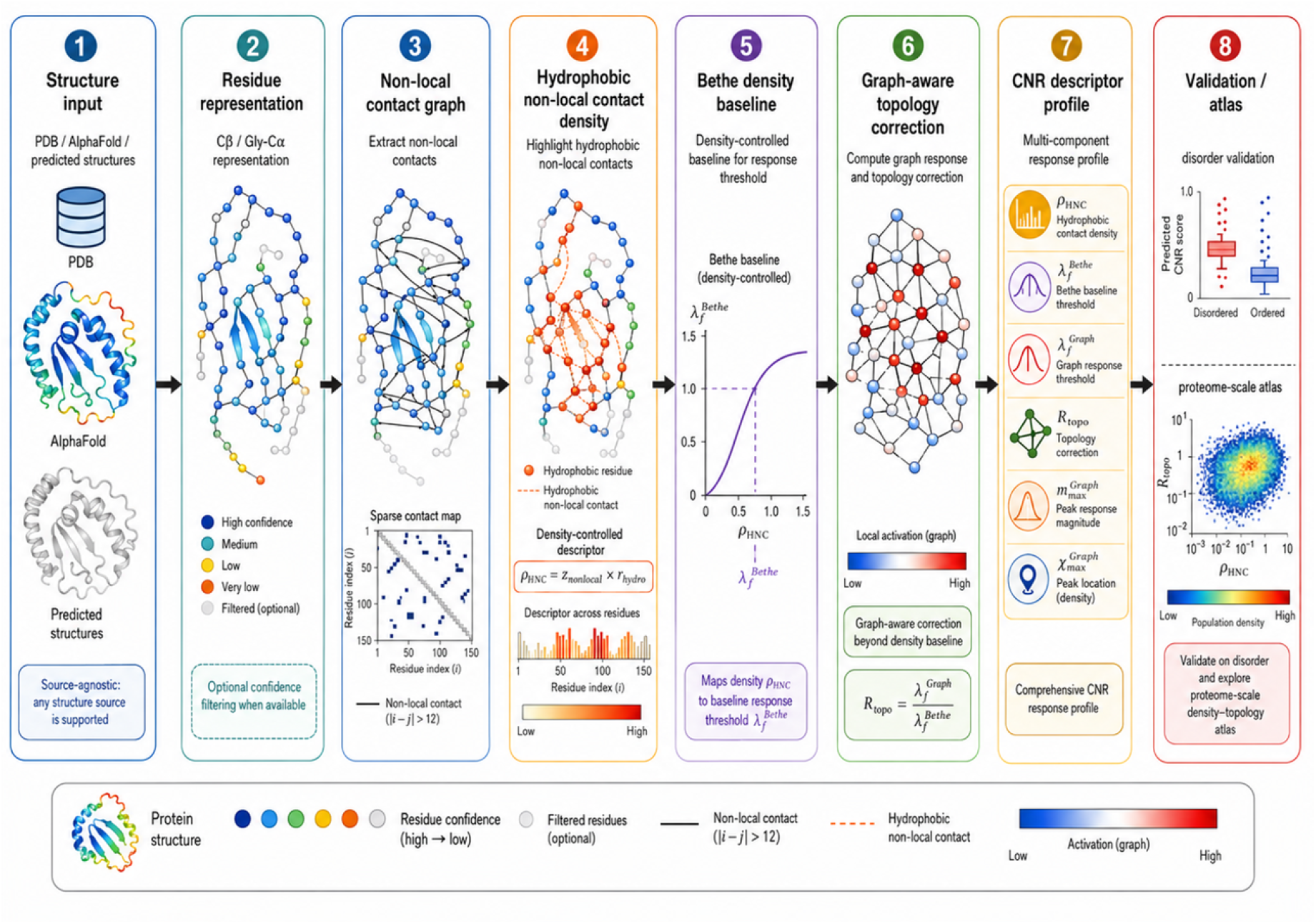
Structure-derived CNR workflow. The CNR framework converts standard protein structures into contact-network responsiveness descriptors. Protein structures from PDB, AlphaFold or other predicted models are first converted into residue-level representations using Cβ atoms, with glycine represented by Cα. Optional confidence filtering can be applied when confidence information is available. Non-local contacts are extracted to form a residue contact graph, and hydrophobic non-local contacts are used to compute the density descriptor *ρ*_HNC_ = *z*_nonlocal_*r*_hydro_. This density descriptor is mapped to a Bethe mean-field baseline response threshold, while the graph-aware layer computes topology correction beyond the density baseline. The final output is a multi-component CNR profile, which can be used for proteome-scale atlas construction and external validation against disorder annotations.

**Figure 2.**
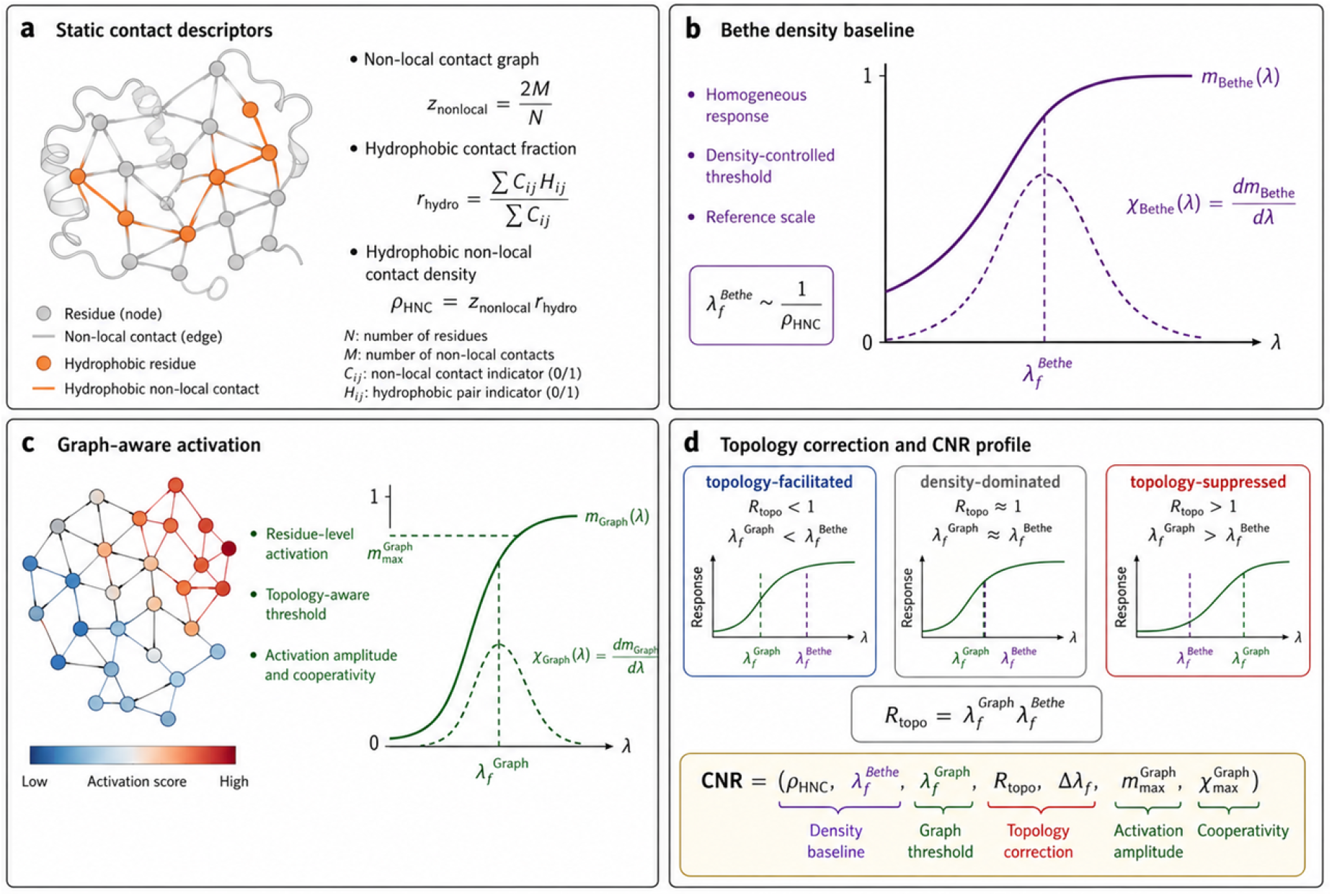
Definition of the CNR descriptor profile. (a)Static contact descriptors extracted from a protein structure. Residues are represented as nodes and non-local contacts as edges. Hydrophobic residues and hydrophobic non-local contacts are used to compute the hydrophobic contact fraction *r*_hydro_, the effective non-local coordination number *z*_nonlocal_, and the hydrophobic non-local contact density *ρ*_HNC_ = *z*_nonlocal_*r*_hydro_. (b) Bethe density baseline. The homogeneous Bethe layer converts *ρ*_HNC_ into a density-controlled response curve *m*_Bethe_(*λ*). The Bethe response threshold 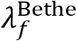 is defined as the point of maximal susceptibility, argmax_*λ*_*dm*_Bethe_/*dλ*, and provides the mean-field reference scale for each protein. (c) Graph-aware residue-level activation. The graph layer uses the actual hydrophobic non-local contact graph to compute residue-level activation variables *m*_*i*_(*λ*), the graph-averaged response *m*_Graph_(*λ*), the graph threshold 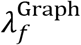, the maximum activation 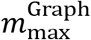, and the maximum susceptibility 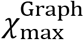. (d) Topology correction and final CNR profile. The topology correction ratio 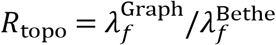 measures whether the actual contact topology facilitates or suppresses activation relative to the density-controlled Bethe baseline. The final CNR profile combines density, threshold, topology-correction, activation-amplitude and susceptibility descriptors.

In this study, CNR was applied to 22,167 valid human protein structures after deduplication and filtering. Median values in the first atlas were *ρ*_HNC_ = 1.43, 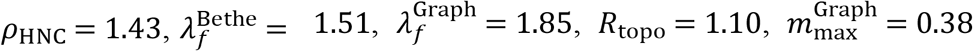, and 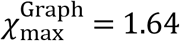. These values provide a proteome-wide baseline for comparing proteins in CNR space.

### Hydrophobic non-local contact density defines the mean-field response baseline

We first asked how much of the response can be explained by average hydrophobic contact density alone. In the Bethe layer, each protein is reduced to *z*_nonlocal_ and *r*_hydro_. The susceptibility threshold is defined as

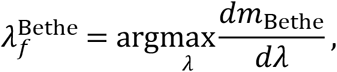

where *m*_Bethe_ is the homogeneous contact-competent occupancy. Across 22,167 valid proteins, 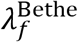 collapsed almost entirely onto *ρ*_HNC_ (Fig. 3a). The relationship was well approximated by

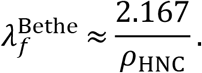

**Figure 3.**
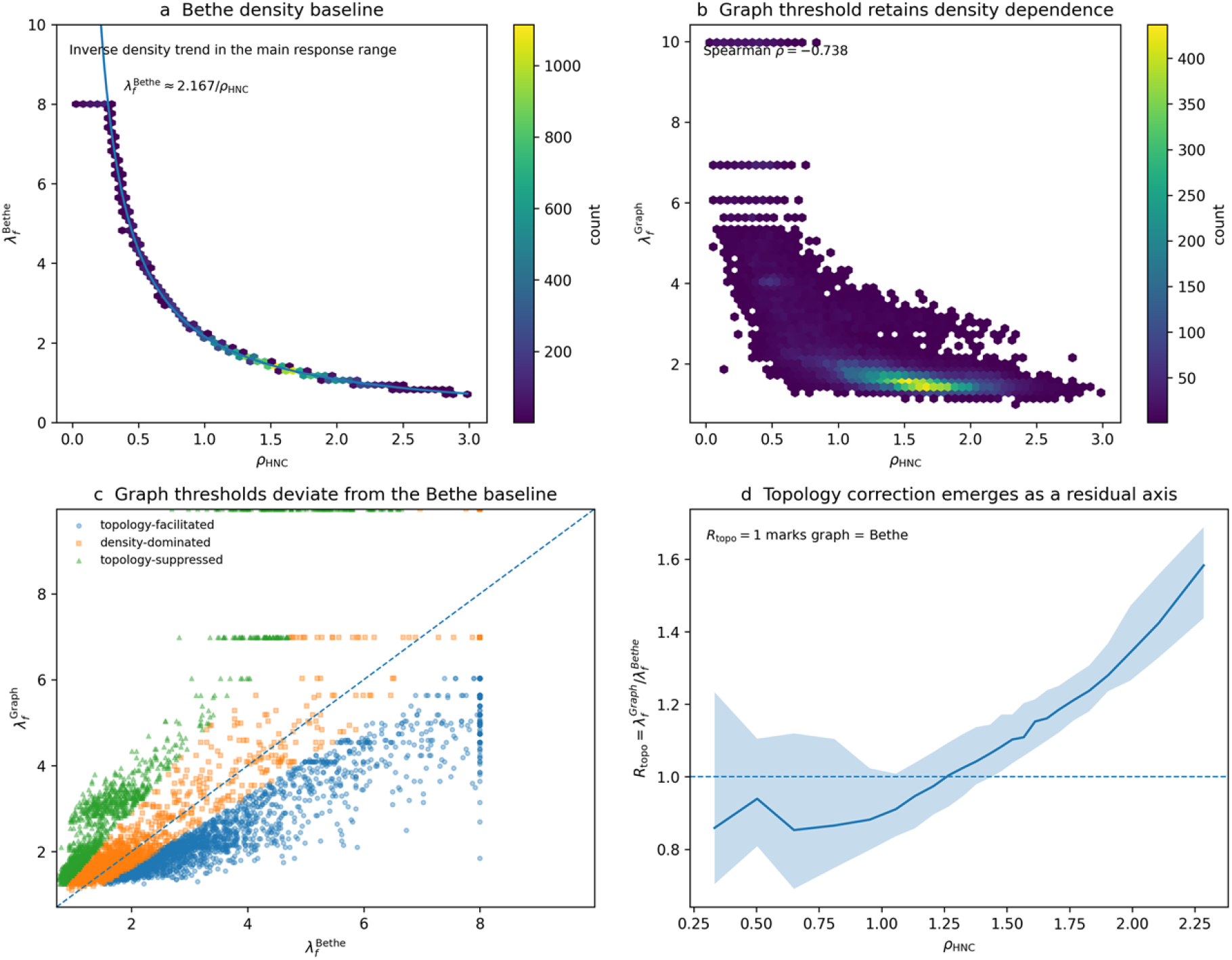
Density baseline and graph-derived topology correction. (a) Bethe response threshold 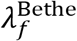 as a function of hydrophobic non-local contact density *ρ*_HNC_. The fitted inverse relation 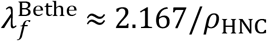 defines the density-controlled mean-field baseline. (b) Graph-aware response threshold 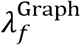 as a function of *ρ*_HNC_. Compared with the Bethe baseline, the graph threshold shows weaker collapse onto contact density, indicating additional topology-dependent variation. (c) Direct comparison of 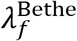 and 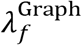, colored by topology-correction class. Deviation from the diagonal indicates graph-derived correction relative to the density baseline. (d) Topology correction ratio 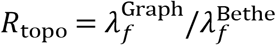 summarized across *ρ*_HNC_ bins. The horizontal line at *R*_topo_ = 1 marks equality between graph and Bethe thresholds. The line indicates the median and the shaded region indicates the interquartile range.

The Spearman correlation between *ρ*_HNC_ and 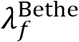 was ― 0.9998. This near-complete collapse is not presented as a universal law of protein folding. It establishes the density baseline of CNR: average hydrophobic non-local contact density sets the first-order response scale in the homogeneous mean-field approximation.

This baseline is essential because it provides the reference needed to interpret topology. Without the Bethe layer, graph thresholds would be difficult to compare across proteins with different contact densities. With the baseline, graph-aware deviations can be quantified as topology correction.

Graph-aware response defines a topology correction beyond the density baseline

The Bethe layer first established a density-controlled reference for CNR. Across 22,167 valid human protein structures, the Bethe response threshold 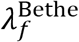 collapsed almost entirely onto hydrophobic non-local contact density, following the inverse relation 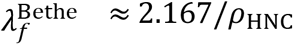 (Fig. 3a). This near-complete collapse indicates that, in the homogeneous mean-field approximation, *ρ*_HNC_ sets the primary response scale.

We then asked whether the actual hydrophobic contact graph provided information beyond this density baseline. The graph-aware threshold 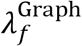 remained associated with *ρ*_HNC_, but the collapse was substantially weaker than for the Bethe threshold (Fig. 3b). Whereas the Spearman correlation between *ρ*_HNC_ and 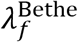 was ― 0.9998, the corresponding correlation for 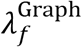 was ― 0.7383. This reduction indicates that the graph-aware response is not a simple restatement of hydrophobic contact density.

Direct comparison of 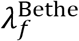 and 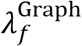 further showed systematic deviation from the density-controlled baseline (Fig. 3c). To quantify this deviation, we defined the topology correction ratio

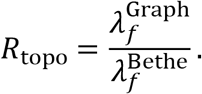

Proteins with *R*_topo_ < 1 have graph thresholds lower than the Bethe baseline and are interpreted as topology-facilitated; proteins with *R*_topo_ ≈ 1 are density-dominated; and proteins with *R*_topo_ > 1 are topology-suppressed. Summarizing *R*_topo_ across *ρ*_HNC_ bins showed that topology correction forms a residual response axis relative to the density baseline, rather than a duplicate density descriptor (Fig. 3d). Thus, Fig. 3 establishes the basic CNR decomposition: hydrophobic contact density defines the mean-field baseline, while graph topology defines the protein-specific correction.

### Atlas-level organization of topology correction in CNR space

To examine how topology correction is organized across the human protein atlas, we mapped all valid proteins into a two-dimensional CNR space defined by hydrophobic non-local contact density and the topology correction ratio. In this representation, *ρ*_HNC_ provides the density-controlled structural baseline, whereas *R*_topo_ measures how the actual hydrophobic contact graph shifts the graph-aware response threshold relative to that baseline. Proteins occupied a continuous density–topology landscape rather than forming sharply separated clusters (Fig. 4a), indicating that topology correction is best interpreted as a graded structural-response axis.

**Figure 4.**
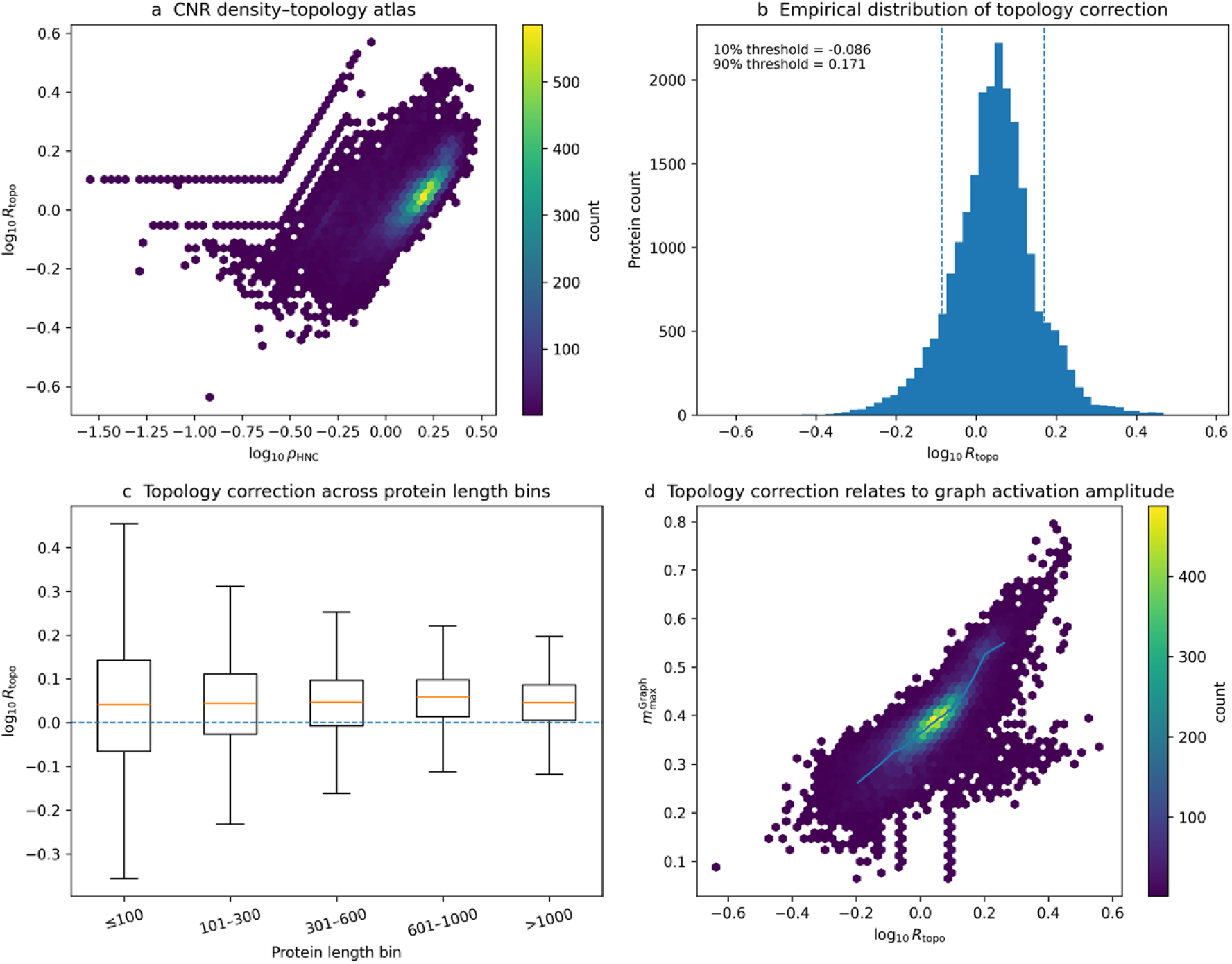
Atlas-level organization of topology correction in CNR space. (a) CNR density–topology atlas showing the joint distribution of hydrophobic non-local contact density *ρ*_HNC_ and topology correction ratio *R*_topo_, shown on log_10_ scales. The horizontal axis represents the density-controlled structural baseline, whereas the vertical axis represents graph-topology correction relative to that baseline. (b) Distribution of log_10_*R*_topo_. Dashed lines mark the lower and upper 10% thresholds used to define topology-facilitated and topology-suppressed proteins. Proteins between these thresholds are classified as density-dominated. (c) Distribution of log_10_*R*_topo_ across protein length bins. Shorter proteins show broader dispersion of topology correction, whereas longer proteins exhibit a more compact distribution. (d) Relationship between log_10_*R*_topo_ and graph activation amplitude 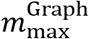. The running median indicates that topology correction is linked to graph-aware activation behavior rather than being merely a classification label.

The distribution of log_10_*R*_topo_ was centered near the density-dominated regime, but extended into facilitated and suppressed tails (Fig. 4b). We therefore defined the lower and upper 10% tails of this empirical distribution as topology-facilitated and topology-suppressed proteins, respectively, while the central 80% were classified as density-dominated. This classification does not impose discrete mechanistic categories a priori; rather, it provides an operational way to identify proteins whose graph-aware response deviates most strongly from the density-controlled baseline.

We next examined whether topology correction was primarily a length-dependent artifact. Across length bins, log_10_*R*_topo_ remained centered near the density-dominated regime, but shorter proteins showed broader dispersion than longer proteins (Fig. 4c). This pattern is consistent with the greater variability of non-local contact organization in small proteins, where a limited number of contacts can produce larger relative deviations from the mean-field baseline. Longer proteins showed a more compact distribution, suggesting that larger contact networks average out local topological fluctuations more effectively.

Finally, topology correction was systematically related to graph activation amplitude. Proteins with different *R*_topo_ values also differed in 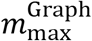, indicating that *R*_topo_ is not merely a bookkeeping ratio between two thresholds (Fig. 4d). Instead, topology correction is linked to the magnitude of graph-aware contact-network activation. These atlas-level analyses show that topology correction is a structured component of CNR: proteins occupy a continuous density–topology space, empirical tails define facilitated and suppressed response regimes, and the topology axis is related to both protein length and graph activation behavior.

### CNR descriptors are not reducible to AlphaFold confidence

Because the present atlas uses AlphaFold models, we tested whether the new CNR descriptors simply reproduce AlphaFold confidence. This is an important control, because low-confidence or disordered regions may influence contact extraction. The topology correction ratio was nearly independent of retained mean pLDDT:

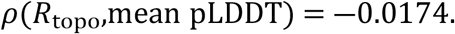

Graph activation showed a weak positive association with the retained residue fraction, but the magnitude was modest. Thus, while confidence filtering affects which residues enter the contact graph, the topology correction descriptor is not a re-labeled AlphaFold confidence metric. This supports the interpretation that CNR captures contact-network organization rather than simply reflecting structural confidence.

### CNR captures a contact-network organization axis associated with intrinsic disorder

We next tested whether CNR descriptors align with independent disorder annotations. Intrinsically disordered proteins and regions are expected to have weaker or less complete non-local contact organization. Therefore, if CNR captures a meaningful contact-network organization axis, higher disorder content should generally be associated with lower hydrophobic contact density, higher response thresholds, and weaker graph activation.

We first used curated DisProt annotations as a high-confidence but limited-coverage benchmark. Across matched human proteins, DisProt disorder content showed directionally consistent associations with CNR descriptors: higher disorder content was associated with lower *ρ*_HNC_, higher 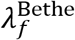, higher 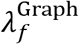, and lower 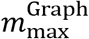. The effect sizes were modest, which is consistent with the curated, heterogeneous and incomplete nature of DisProt coverage. These results were used as an initial external check rather than as the primary validation.

We then performed a broader validation using UniProt region annotations derived from MobiDB-lite. Disordered regions were parsed from UniProt Region entries annotated as “Disordered”, and a disorder fraction was computed for each protein. Across 22,020 matched CNR protein models, predicted disorder fraction was systematically associated with the CNR profile. Disorder fraction was negatively correlated with hydrophobic non-local contact density (*ρ* = ― 0.288), positively correlated with the Bethe response threshold (*ρ* = 0.289), positively correlated with the graph response threshold (*ρ* = 0.195), negatively correlated with graph maximum activation (*ρ* = ― 0.244), and negatively correlated with graph susceptibility peak (*ρ* = ― 0.215).

The trend was visible both as continuous correlations (Fig. 6) and by disorder class (Fig. 5). Proteins without annotated disordered regions had a median *ρ*_HNC_ of 1.546, median 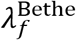 of 1.395, and median 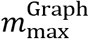 of 0.395. Proteins with very high disorder content had a median *ρ*_HNC_ of 0.682, median 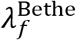 of 3.205, and median 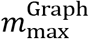 of 0.296. Thus, as disorder content increased, hydrophobic contact density decreased, response thresholds increased, and graph activation weakened.

**Figure 5.**
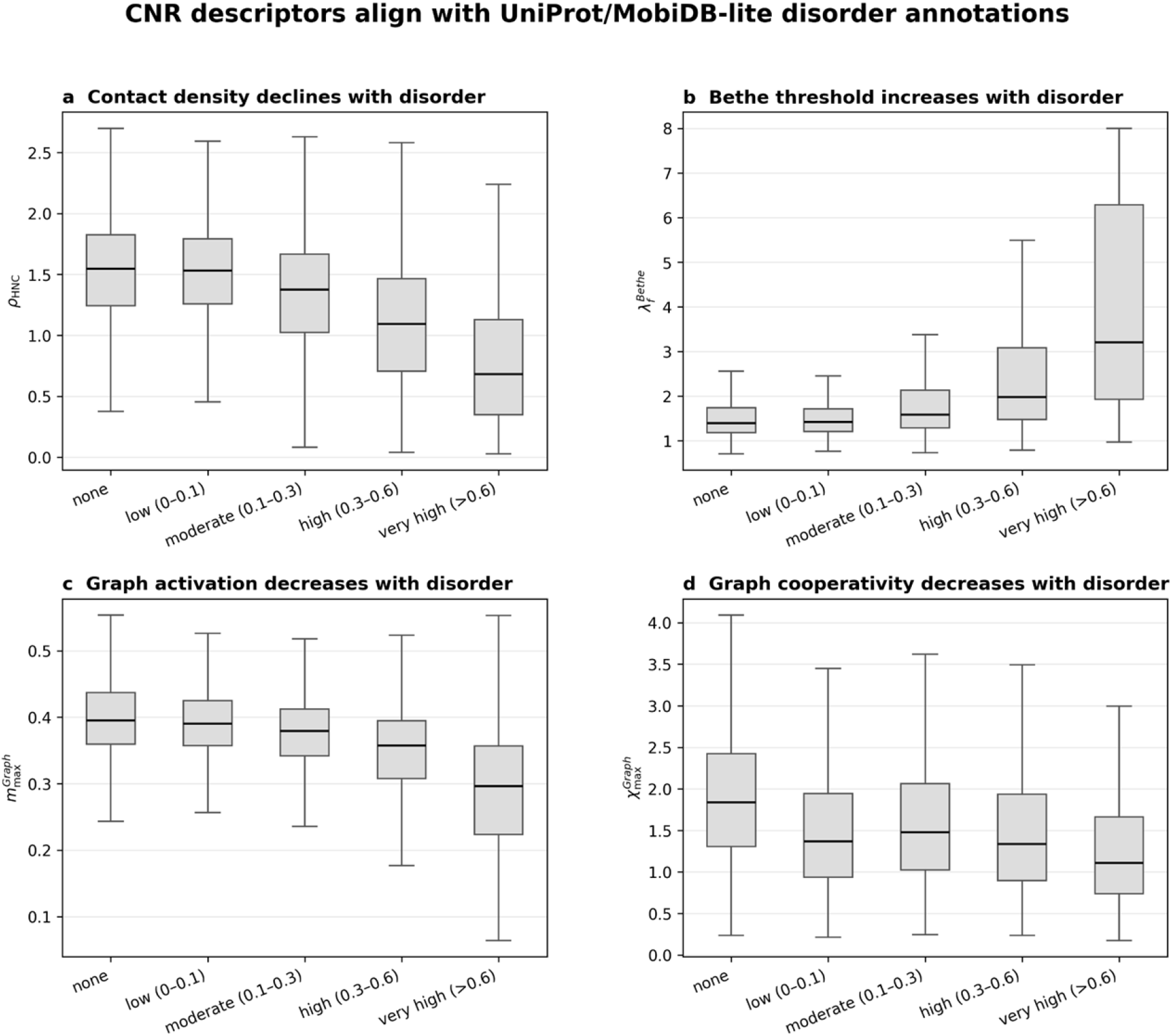
CNR descriptors across UniProt/MobiDB-lite disorder classes. (a) Hydrophobic non-local contact density across disorder classes. (b) Bethe response threshold across disorder classes. (c) Graph maximum activation across disorder classes. (d) Graph susceptibility peak across disorder classes. Higher disorder content is associated with lower contact density, higher response thresholds and weaker graph activation.

**Figure 6.**
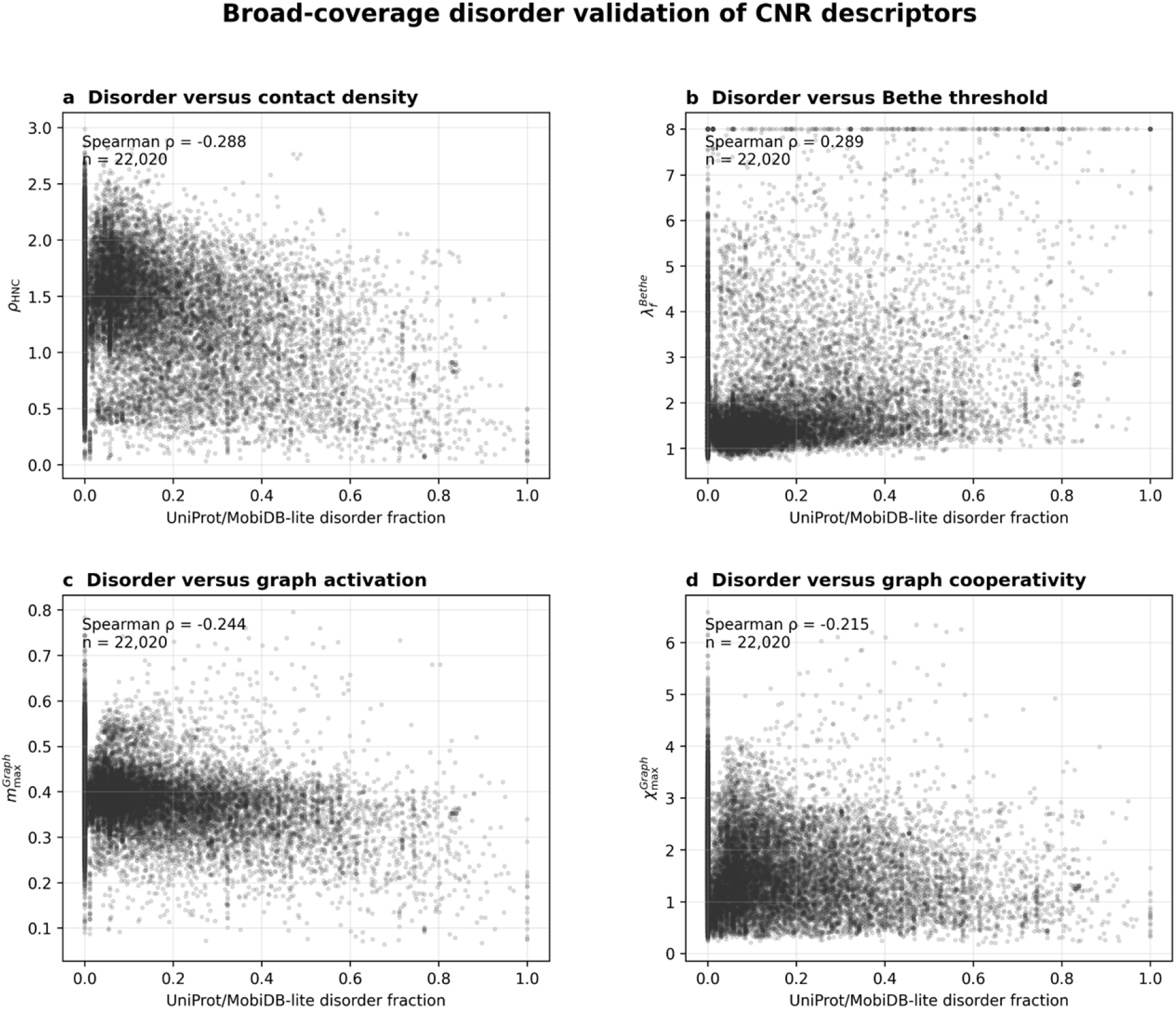
Continuous disorder-fraction validation of CNR descriptors. (a) UniProt/MobiDB-lite disorder fraction versus hydrophobic non-local contact density. (b) Disorder fraction versus Bethe response threshold. (c) Disorder fraction versus graph maximum activation. (d) Disorder fraction versus graph susceptibility peak. These panels provide the continuous-form validation corresponding to the disorder-class trends in Fig. 5.

These results do not mean that CNR is a disorder predictor. Instead, disorder provides an external benchmark for low-density and low-activation CNR regimes. The associations support the interpretation that CNR captures a contact-network organization axis related to the availability and responsiveness of non-local hydrophobic contacts.

### Representative protein cases illustrate distinct CNR regimes

To make the CNR descriptors interpretable at the single-protein level, we selected four representative proteins from distinct regions of the CNR atlas (Fig. 7). These examples were not used to define the model, but were chosen to illustrate how density, topology correction, activation amplitude and cooperativity combine into distinct response profiles.

**Figure 7.**
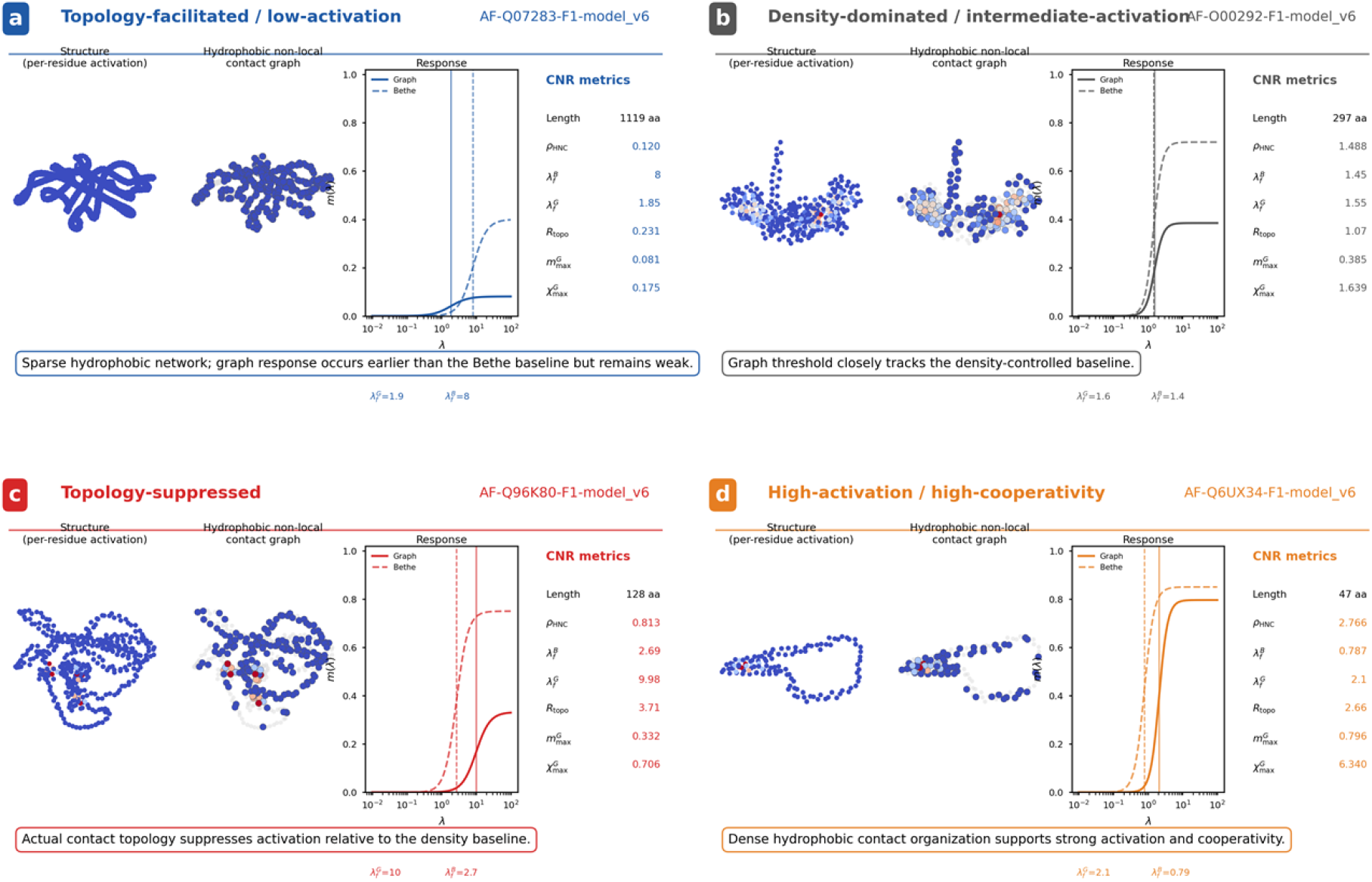
Representative CNR protein cases from actual structure files. Four representative proteins illustrate distinct regions of the CNR atlas. Structure panels and hydrophobic non-local contact graphs were rebuilt from the corresponding AlphaFold structure files using the same residue-representation logic used for CNR analysis: Cβ atoms were used for all residues except glycine, for which Cα was used. Non-local contacts were defined with an 8 Å distance cutoff after excluding contacts between residues separated by 12 or fewer sequence positions; hydrophobic non-local contacts are shown in the graph panels. Response-curve insets summarize the computed CNR thresholds and amplitudes for each representative protein. (a) AF-Q07283-F1-model_v6 represents a topology-facilitated but low-activation regime. (b) AF-O00292-F1-model_v6 represents a density-dominated intermediate-activation regime. (c) AF-Q96K80-F1-model_v6 represents a topology-suppressed regime. (d) AF-Q6UX34-F1-model_v6 represents a high-activation, high-cooperativity regime.

The topology-facilitated low-activation case, AF-Q07283-F1-model_v6, had a very low hydrophobic non-local contact density (*ρ*_HNC_ = 0.120), a high Bethe threshold 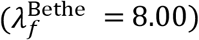, and an earlier graph threshold 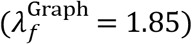, yielding *R*_topo_ = 0.231. Despite this early graph response, the maximum graph activation was low 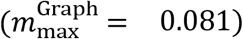. This case illustrates that topology facilitation does not necessarily imply strong global activation; sparse contact networks can respond earlier than the density baseline while remaining weak in amplitude.

The density-dominated case, AF-O00292-F1-model_v6, showed close agreement between the Bethe and graph thresholds 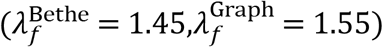, with *R*_topo_ = 1.07. Its intermediate graph activation 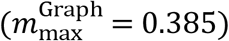 and susceptibility peak 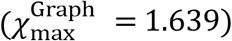 are consistent with a regime in which the graph-aware response closely follows the density-controlled baseline.

The topology-suppressed case, AF-Q96K80-F1-model_v6, had *ρ*_HNC_ = 0.813, but the graph threshold was strongly delayed relative to the Bethe baseline 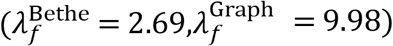, yielding *R*_topo_ = 3.71. This example illustrates a protein for which the average density descriptor alone would underestimate the effective response threshold because the actual graph topology suppresses activation.

Finally, the high-activation/high-cooperativity case, AF-Q6UX34-F1-model_v6, had high hydrophobic contact density (*ρ*_HNC_ = 2.766), high graph activation 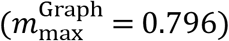, and a large graph susceptibility peak 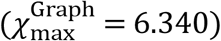. This case illustrates that dense hydrophobic contact organization can support a strong, cooperative graph response. Together, these examples illustrate how the CNR profile should be interpreted: density sets the baseline, topology shifts the threshold, and graph activation descriptors report the amplitude and sharpness of contact-network response.

## Discussion

This work introduces CNR as a structure-derived framework for contact-network responsiveness. Its central purpose is to convert static protein coordinates into interpretable response descriptors. Unlike traditional contact-density measures, CNR separates a density-controlled baseline from topology-dependent correction. Unlike molecular dynamics, it does not attempt to sample explicit conformational ensembles. Instead, it provides a scalable annotation layer that can be applied to large structural datasets.

The first layer of CNR is the density baseline. The near-exact relationship between 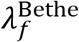 and *ρ*_HNC_ shows that hydrophobic non-local contact density controls the homogeneous mean-field response threshold. This finding is important because it establishes a transparent reference scale. The Bethe baseline is not intended as a complete folding theory; it is the baseline against which graph topology can be measured.

The second layer is topology correction. The graph-aware model incorporates the actual residue contact graph, and *R*_topo_ converts deviation from the Bethe baseline into a protein-level descriptor. This allows proteins to be classified as topology-facilitated, density-dominated, or topology-suppressed. In this sense, CNR asks a question not answered by static contact density alone: given the same average hydrophobic contact density, does the actual contact topology make network activation easier or harder?

The third layer is biological validation. Both curated DisProt annotations and broad-coverage UniProt/MobiDB-lite disorder annotations support the interpretation that CNR tracks a contact-network organization axis. Higher disorder content is associated with lower hydrophobic contact density, higher response thresholds and weaker graph activation. These relationships are not perfect and should not be overinterpreted. Intrinsic disorder is only one biological manifestation of weak or incomplete contact organization. Nevertheless, the disorder validation demonstrates that CNR descriptors are not merely internally consistent mathematical outputs; they relate to an independent structural-biological annotation.

The framework is source-agnostic. Although the present atlas uses human AlphaFold structures, CNR requires only residue-level coordinates and can be applied to experimentally determined PDB structures, predicted models, homologous protein families, mutation series, or alternative conformational states. This generality is important. The term “AlphaFold-derived” describes the substrate used in this study, not the theoretical scope of the method.

CNR also has practical value as a first-pass annotation resource. The descriptor table can be used to rank proteins by hydrophobic contact density, topology correction, activation amplitude, or cooperativity before selecting targets for more expensive molecular simulation or experimental validation. In this role, CNR is best understood as a screening and annotation layer rather than a final thermodynamic measurement.

Several limitations should be emphasized. First, CNR does not predict experimental melting temperature, folding free energy, or kinetic folding rate. Second, the current hydrophobic interaction representation is simplified and does not explicitly model electrostatics, hydrogen bonding, disulfide bonds, metal coordination, ligand effects or solvent-mediated interactions. Third, the graph-aware model is still mean-field: residue occupancies are self-consistent activation variables, not directly measured folded fractions. Fourth, disorder annotations from UniProt/MobiDB-lite are prediction-derived, although DisProt provides a curated complementary benchmark.

Future work can improve CNR in several directions. Residue-pair potentials, electrostatic terms, hydrogen-bond propensities, secondary-structure context and confidence-weighted contacts could be incorporated into the contact weights. Mutation analysis can be implemented by locally modifying residue-contact weights and recalculating the CNR profile. Experimentally, selected proteins from topology-facilitated, topology-suppressed and low-activation regimes could be tested using differential scanning fluorimetry, circular dichroism, limited proteolysis or aggregation assays. Finally, residue-level graph activation can be mapped back onto structures to identify regions predicted to activate early or late as effective contact strength increases.

In summary, CNR provides a structure-derived route from static protein coordinates to contact-network responsiveness. Hydrophobic non-local contact density defines the mean-field baseline; graph topology defines the correction; and disorder annotations provide an external benchmark for weak contact-network organization. This structure-source-agnostic framework may serve as a first-pass annotation layer for proteome-scale structural datasets and for future studies of mutation, disorder, stability and contact-network organization.

## Methods

### Structure dataset

Human AlphaFold protein structures were used as the high-coverage structural substrate for this atlas [4,5]. Duplicate structure formats were removed. When both .cif.gz and .pdb.gz versions were present for the same model identifier, the .cif.gz file was retained. After deduplication and filtering, 22,167 protein models passed the contact and model-validity criteria. Although AlphaFold structures were used in this study, the CNR framework itself requires only residue-level coordinates and can be applied to experimentally determined or predicted protein structures [4–6].

### Contact extraction

Residues were represented by Cβ atoms, except glycine, which was represented by Cα. AlphaFold pLDDT values were read from the B-factor field, and low-confidence residues were excluded before contact extraction. Non-local contacts were defined using an 8 Å cutoff between residue representatives and a sequence-separation exclusion. Peptide-bond connectivity was treated as a fixed chain constraint and was not counted as a stabilizing non-local contact.

For a protein with *N* retained residues and *M* undirected non-local contacts, the effective non-local coordination number was

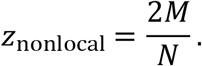

Hydrophobic contacts were defined using the Kyte-Doolittle scale, with positive-scale residues classified as hydrophobic [12]. The hydrophobic contact fraction was

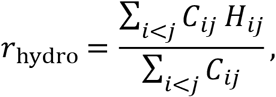

where *H*_*ij*_ = 1 for hydrophobic-hydrophobic contacts and 0 otherwise. The hydrophobic non-local contact density was *ρ*_HNC_ = *z*_nonlocal_*r*_hydro_.

### Bethe baseline

For each valid protein, a q-state Bethe mean-field model was solved using *q* = 3, *z*_nonlocal_ and *r*_hydro_ [13]. The cavity fields obeyed

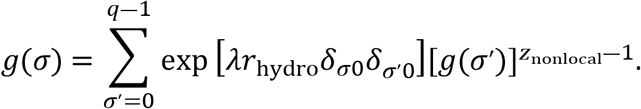

After convergence, the homogeneous contact-competent occupancy was

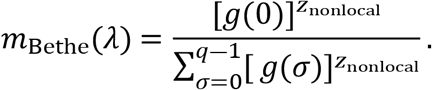

The Bethe response was scanned over 0.1 ≤ *λ* ≤ 8.0 using 300 grid points. The susceptibility threshold 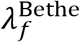 was defined as the grid point where *dm*_Bethe_/*dλ* was maximal.

### Graph-aware model

The graph-aware model used the hydrophobic non-local contact graph [1,14]. Residue occupancies were solved from

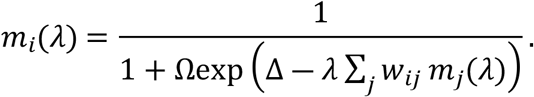

The calibrated analysis used Δ = 2.5 and Ω = 2.0. The graph response was scanned over 0.1 ≤ *λ* ≤ 20.0 using 400 grid points. Graph-level occupancy was *m*_Graph_ = *N*^―1^∑_*i*_ *m*_*i*_. The graph threshold, maximum activation and maximum susceptibility were computed from the graph response curve.

### Topology classes

The topology correction ratio was defined as

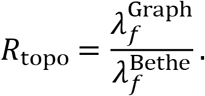

Proteins in the lower 10% tail of log_10_*R*_topo_ were classified as topology-facilitated, proteins in the upper 10% tail were classified as topology-suppressed, and all remaining proteins were classified as density-dominated.

### Disorder validation

Curated disorder annotations were obtained from DisProt and merged with CNR descriptors using UniProt accession identifiers [7–9]. For broad-coverage validation, UniProt reviewed human protein entries were downloaded with Region annotations [7,10,11]. Disordered regions were parsed from entries annotated as /note=“Disordered”, which commonly included MobiDB-lite evidence. Disorder lengths were merged across overlapping intervals and divided by sequence length to compute per-protein disorder fraction.

Associations between CNR descriptors and disorder fraction were quantified using Spearman rank correlation. Proteins were grouped into disorder classes: none, low, moderate, high and very high disorder content.

### Statistical analysis

Distributions were summarized using medians, quartiles, means and standard deviations. Associations between continuous variables were quantified using Spearman rank correlation. Protein length was grouped into predefined bins. Proteins with too few retained residues, insufficient non-local contacts or zero hydrophobic contact weight were excluded from model-valid response statistics.

## Data availability

The CNR descriptor tables, merged disorder-validation tables and figure source data are available upon request. The input structures used for the human proteome-scale atlas were obtained from the AlphaFold Protein Structure Database. Curated intrinsic-disorder annotations were obtained from DisProt, and broad-coverage disorder annotations were parsed from UniProt reviewed human protein Region annotations containing MobiDB-lite-supported disordered regions. The four structure files used for representative case visualization are listed by AlphaFold model identifier in Fig. 7.

## Code availability

The CNR calculation scripts used to extract non-local contacts, compute Bethe baseline descriptors, compute graph-aware response descriptors and generate the atlas tables are deposited at: https://github.com/ranhuang84/CNR-Structure-Atlas.

## Acknowledgements

The work is financially supported by the YRI-FD Industrial Project (YRI-IP-25-01).

## CRediT authorship contribution statement

**Ran Huang**: Conceptualization, Data curation, Formal analysis, Investigation, Resources, Methodology, Software, Validation, Writing - original draft, Writing – review & editing. **Xin Ma**: Conceptualization, Funding acquisition, Resources, Project administration, Supervision, Writing - review & editing. **Dean Ta:** Conceptualization, Funding acquisition, Resources, Project administration, Supervision, Writing - review & editing.

## Competing interests

The authors declare no competing interests.

## AI-assisted preparation statement

AI-assisted tools were used to support language editing, code drafting and schematic figure prototyping. Scientific interpretation, data selection, final analyses and manuscript approval remain the responsibility of the authors.

## Notes

### Competing Interest Statement

The authors have declared no competing interest.

